# Family-wide evaluation of RALF peptides in *Arabidopsis thaliana*

**DOI:** 10.1101/2020.06.26.174169

**Authors:** Alicia Abarca, Christina M. Franck, Cyril Zipfel

## Abstract

Plant peptide hormones are important players controlling various aspects of plants’ lives. RAPID ALKALINIZATION FACTOR (RALF) peptides have recently emerged as important players in multiple physiological processes. Numerous studies on RALF peptides focused on broad phylogenetic analysis including multiple species. Thus, progress has been made in understanding the evolutionary processes that shaped this family. Nevertheless, to date, there is no comprehensive, family-wide functional study on RALF peptides. Here, we analysed the phylogeny and function of the proposed multigenic RALF peptide family in the model plant *Arabidopsis thaliana*, ecotype Col-0. Our phylogenetic analysis reveals that two of the previously proposed RALF peptides are not genuine RALF peptides, which leads us to propose a new consensus *At*RALF peptide family annotation. Moreover, we show that the majority of *At*RALF peptides are able to induce seedling or root growth inhibition in *A. thaliana* seedlings when applied exogenously as synthetic peptides. Additionally, we show that most of these responses are dependent on the *Catharanthus roseus* RLK1-LIKE receptor kinase FERONIA, suggesting a pivotal role in the perception of multiple RALF peptides.

**One sentence summary:** The majority of *A. thaliana* RALF peptides inhibit growth in a FERONIA-dependent manner

## Introduction

Cell-to-cell communication is crucial for plants, as sessile organisms constantly exposed to an ever-changing environment. Under this scenario, plant peptide hormones are key to rapidly initiate, coordinate and integrate responses thanks to their large diversity in structure, function and expression patterns (Matsubayashi, 2014; Olsson et al., 2019).

RAPID ALKALINIZATION FACTOR (RALF) peptides belong to a family of cysteine-rich plant peptide hormones that are involved in multiple physiological and developmental processes, ranging from pollen tube growth to modulation of immune responses (Murphy & De Smet, 2014; Blackburn et al., 2020). They were discovered in a peptide hormone screen due to their ability to cause medium alkalinisation of tobacco cell cultures (Pearce et al., 2001a). Later, RALF peptides, with several conserved motifs, were found to be ubiquitous in terrestrial plants highlighting their functional importance (Cao & Shi, 2012; Murphy & De Smet, 2014; Campbell & Turner, 2017).

In *Arabidopsis thaliana*, more than 30 RALF peptides have been predicted. In the ecotype Col-0, various studies have reported between 34 and 39 members, depending on the study and criteria considered (Olsen et al., 2002; Cao & Shi, 2012; Haruta et al., 2014; Morato do Canto et al., 2014; Sharma et al., 2016; Stegmann et al., 2017; Campbell & Turner, 2017). This discrepancy calls for a careful examination of the RALF family annotation.

The majority of plant peptide hormones are perceived by plasma-membrane localized leucine-rich repeat receptor kinases (LRR-RKs) or receptor proteins (LRR-RPs), which unlike RKs lack an intracellular domain (Hohmann et al., 2017; Olsson et al., 2019). In contrast, RALF peptides have recently been shown to be ligands of protein complexes involving *Catharanthus roseus* RLK1-LIKE (CrRLK1L) receptor kinases, characterized by malectin-like domains in their extracellular domain (Franck et al., 2018). For example, RALF1, RALF22 and RALF23 bind to the CrRLK1L FERONIA (FER) to regulate root growth, abiotic, and biotic stress responses, respectively (Haruta et al., 2014; Stegmann et al., 2017; Zhao et al., 2018). RALF4 and RALF19 were shown to be ligands for the CrRLK1Ls ANXUR1 (ANX1), ANX2 and BUDDHA’S PAPER SEAL (BUPS) 1 and BUPS2 in the context of pollen tube growth and integrity (Ge et al., 2017). RALF34 was recently shown to bind the CrRLK1L THESEUS1 (THE1) to regulate growth upon cellulose biosynthesis inhibition (Gonneau et al., 2018). CrRLK1Ls, such as FER, ANX1/ANX2 and BUPS1/2, have been shown to work together with the glycosylphosphatidylinositol-anchored protein (GPI-AP) LORELEI or related (LRE)-like-GPI-anchored proteins (LLGs) to perceive RALF peptides (Xiao et al., 2019; Ge et al., 2019; Feng et al., 2019). Notably, RALF peptides can also bind LEUCINE-RICH REPEAT EXTENSINS (LRX) proteins (Mecchia et al., 2017; Zhao et al., 2018; Moussu et al., 2020). The biochemical relationship between RALF perception by CrRLK1L/LLG complexes and LRXs is still unclear.

There are 17 CrRLK1Ls, 11 LRXs and 4 LRE/LLGs in *A. thaliana* (Li et al., 2015; Franck et al., 2018; Herger et al., 2020). As such, the diversity of potential assembly modules of the RALF-perception/signalling axis could explain the functional plasticity of this family of peptides. For instance, different RALF peptides might be secreted upon diverse stimuli in different tissues and this, in turn, would trigger the formation of receptor complexes with a combination of the above-mentioned assembly modules. There are numerous studies analysing individual aspects of this complex signalling network (Blackburn et al., 2020). However, the functional role of the majority of RALF peptides and their bioactivity is largely unknown. In this study, we performed a family-wide analysis of *At*RALF peptides using seedling and root growth inhibition as read-outs. In addition to defining the core *A. thaliana* Col-0 RALF family, we show that FER is required for full responsiveness to multiple RALFs, suggesting a pivotal role of this receptor in potentially multiple RALF sensory complexes.

## Results

### Re-annotation of the *Arabidopsis thaliana* Col-0 RALF family

The *At*RALF peptide family consists of more than 30 members, ranging from 34 to 39, depending on the studies considered (Blackburn et al., 2020). In order to define the core *At*RALF family in the Col-0 ecotype, we compared the *At*RALF peptide annotation used in different publications and those available in the TAIR10 and UniProt databases (Table I). Notably, we found some inconsistency both in the number and identity proteins that comprise the family.

**Table I.** Re-annotation of *At*RALFs. Literature comparison of the *At*RALF peptide annotation used in different publications (listed in chronological order) and databases. Pink rows indicate inconsistency in the annotation and the final column coloured un green shows our proposed consensus *At*RALF annotation.

| ATG number | TAIR | Protein accession | Olsen et al., 2002 | Cao and Shi et al., 2012 | Haruta et al., 2014 | Morato do Canto et al., 2014 | Stegmann et al., 2017 | Campbell & Turner et al., 2017 | Consensus |
| --- | --- | --- | --- | --- | --- | --- | --- | --- | --- |
| AT1G02900 | RALF1 | Q9SRY3 | RALF1 | AT1G02900 | RALF1 | RALF1 | RALF1 | AT1G02900 | RALF1 |
| AT1G23145 | RALF2 | A8MQ92 | RALF2 | AT1G23145 | RALF2 | RALF2 | RALF2 | AT1G23145 | RALF2 |
| AT1G23147 | RALF3 | A7REE5 | RALF3 | AT1G23147 | RALF3 | RALF3 | RALF3 | AT1G23147 | RALF3 |
| AT1G28270 | RALF4 | Q6TF26 | RALF4 | AT1G28270 | RALF4 | RALF4 | RALF4 | AT1G28270 | RALF4 |
| AT1G35467 | RALF5 | A8MQI8 | RALF5 | AT1G35467 | RALF5 | RALF5 | RALF5 | AT1G35467 | RALF5 |
| AT1G60625 | RALF6 | A8MQM2 | RALF6 | AT1G60625 | RALF6 | RALF6 | RALF6 | AT1G60625 | RALF6 |
| AT1G60815 | RALF7 | A8MRD4 | RALF7 | AT1G60815 | RALF7 | RALF7 | RALF7 | AT1G60815 | RALF7 |
| AT1G61563 | RALF8 | Q1ECR9 | RALF8 | AT1G61563 | RALF8 | RALF8 | RALF8 | AT1G61563 | RALF8 |
| AT1G61566 | RALF9 | Q3ECL0 | RALF9 | AT1G61566 | RALF9 | RALF9 | RALF9 | AT1G61566 | RALF9 |
| AT2G19020 | RALF10 | O65919 | RALF10 | AT2G19020 | RALF10 | RALF10 | RALF10 | AT2G19020 | RALF10 |
| AT2G19030 | RALF11 | O64466 | RALF11 | AT2G19030 | RALF11 | RALF11 | RALF11 | AT2G19030 | RALF11 |
| AT2G19040 | RALF12 | F4ISE1 | RALF12 | AT2G19040 | RALF12 | RALF12 | RALF12 | AT2G19040 | RALF12 |
| AT2G19045 | RALF13 | F4ISE2 | RALF13 | AT2G19045 | RALF13 | RALF13 | RALF13 | AT2G19045 | RALF13 |
| AT2G20660 | RALF14 | Q9SIU6 | RALF14 | AT2G20660 | RALF14 | RALF14 | RALF14 | AT2G20660 | RALF14 |
| AT2G22055 | RALF15 | A8MQM7 | RALF15 | AT2G22055 | RALF15 | RALF15 | RALF15 | AT2G22055 | RALF15 |
| AT2G32835 | RALF16 | A8MRM1 | RALF16 | AT2G32835 | RALF16 | RALF16 | RALF16 | AT2G32835 | RALF16 |
| AT2G32885 | RALF36 | A8MR00 | - | AT2G32885 | - | RALF17 | RALF17 | AT2G32885 | RALF17 |
| AT2G32890 | RALF17 | O48776 | RALF17 | - | RALF17 | - | - | AT2G32890 | AT2G32890 |
| AT2G33130 | RALF18 | O49320 | RALF18 | AT2G33130 | RALF18 | RALF18 | RALF18 | AT2G33130 | RALF18 |
| AT2G33775 | RALF19 | Q6NME6 | RALF19 | AT2G33775 | RALF19 | RALF19 | RALF19 | AT2G33775 | RALF19 |
| AT2G34825 | RALF20 | A8MQL7 | RALF20 | AT2G34825 | RALF20 | RALF20 | RALF20 | AT2G34825 | RALF20 |
| AT3G04735 | RALF21 | A8MRF9 | RALF21 | AT3G04735 | RALF21 | RALF21 | RALF21 | AT3G04735 | RALF21 |
| AT3G05490 | RALF22 | Q9MA62 | RALF22 | AT3G05490 | RALF22 | RALF22 | RALF22 | AT3G05490 | RALF22 |
| AT3G16570 | RALF23 | Q9LUS7 | RALF23 | AT3G16570 | RALF23 | RALF23 | RALF23 | AT3G16570 | RALF23 |
| AT3G23805 | RALF24 | Q9LK37 | RALF24 | AT3G23805 | RALF24 | RALF24 | RALF24 | AT3G23805 | RALF24 |
| AT3G25165 | RALF25 | Q9LSG0 | RALF25 | AT3G25165 | RALF25 | RALF25 | RALF25 | AT3G25165 | RALF25 |
| AT3G25170 | RALF26 | Q0V822 | RALF26 | AT3G25170 | RALF26 | RALF26 | RALF26 | AT3G25170 | RALF26 |
| AT3G29780 | RALF27 | Q9LH43 | RALF27 | - | RALF27 | RALF27 | RALF27 | AT3G29780 | RALF27 |
| AT4G11510 | RALF28 | Q9LDU1 | RALF28 | AT4G11510 | RALF28 | RALF28 | RALF28 | AT4G11510 | RALF28 |
| AT4G11653 | RALF29 | A8MQP2 | RALF29 | AT4G11653 | RALF29 | RALF29 | RALF29 | AT4G11653 | RALF29 |
| AT4G13075 | RALF30 | A7REH2 | RALF30 | AT4G13075 | RALF30 | RALF30 | RALF30 | AT4G13075 | RALF30 |
| AT4G13950 | RALF31 | Q2HIM9 | RALF31 | AT4G13950 | RALF31 | RALF31 | RALF31 | AT3G16570 | RALF31 |
| AT4G14010 | RALF32 | O23262 | RALF32 | AT4G14010 | RALF32 | RALF32 | RALF32 | AT4G14010 | RALF32 |
| AT4G15800 | RALF33 | Q8L9P8 | RALF33 | AT4G15800 | RALF33 | RALF33 | RALF33 | AT4G15800 | RALF33 |
| AT5G67070 | RALF34 | Q9FHA6 | RALF34 | AT5G67070 | RALF34 | RALF34 | RALF34 | AT5G67070 | RALF34 |
| AT1G60913 | - | A8MRK3 | - | - | - | RALF35 | RALF35 | AT1G60913 | RALF35 |
| AT4G14020 | - | O23263 | - | AT4G14020 | RALF35 | - | - | AT4G14020 | AT4G14020 |
| AT2G32785 | - | A8MR74 | - | - | - | RALF36 | RALF36 | - | RALF36 |
| AT2G32788 | - | A8MRN0 | - | - | - | RALF37 | RALF37 | - | RALF37 |

Despite having low level of amino acid sequence similarity, RALF peptides, have diverse conserved motifs, which have been shown to be important for diverse functions (Pearce et al., 2001a; Matos et al., 2008; Srivastava et al., 2009; Pearce et al., 2010; Stegmann et al., 2017). For example, the dibasic site RR in the canonical AtS1P recognition site which is important for processing of the peptide precursor (Fig 1A, Matos et al., 2008; Srivastava et al., 2009). The N-terminal YI/LSY motif of the mature peptide which was shown to be important for binding to the receptor and alkalinisation activity (residues 112 to 115, Fig 1A, Pearce et al., 2010; Xiao et al., 2019). Additionally, there are other parts of the sequences that show high conservation, such as the C-terminal RCRR(S) motif or the four di-sulfide bond-forming cysteine residues that are important for the bioactivity of the peptide (residues 164 to 167, 130, 143, 159 and 165, respectively, Fig 1A, Pearce et al., 2001b; Haruta et al., 2008). Using these criteria, we re-analysed the published RALF family members and propose the following updates to the annotation.

**Fig. 1.**
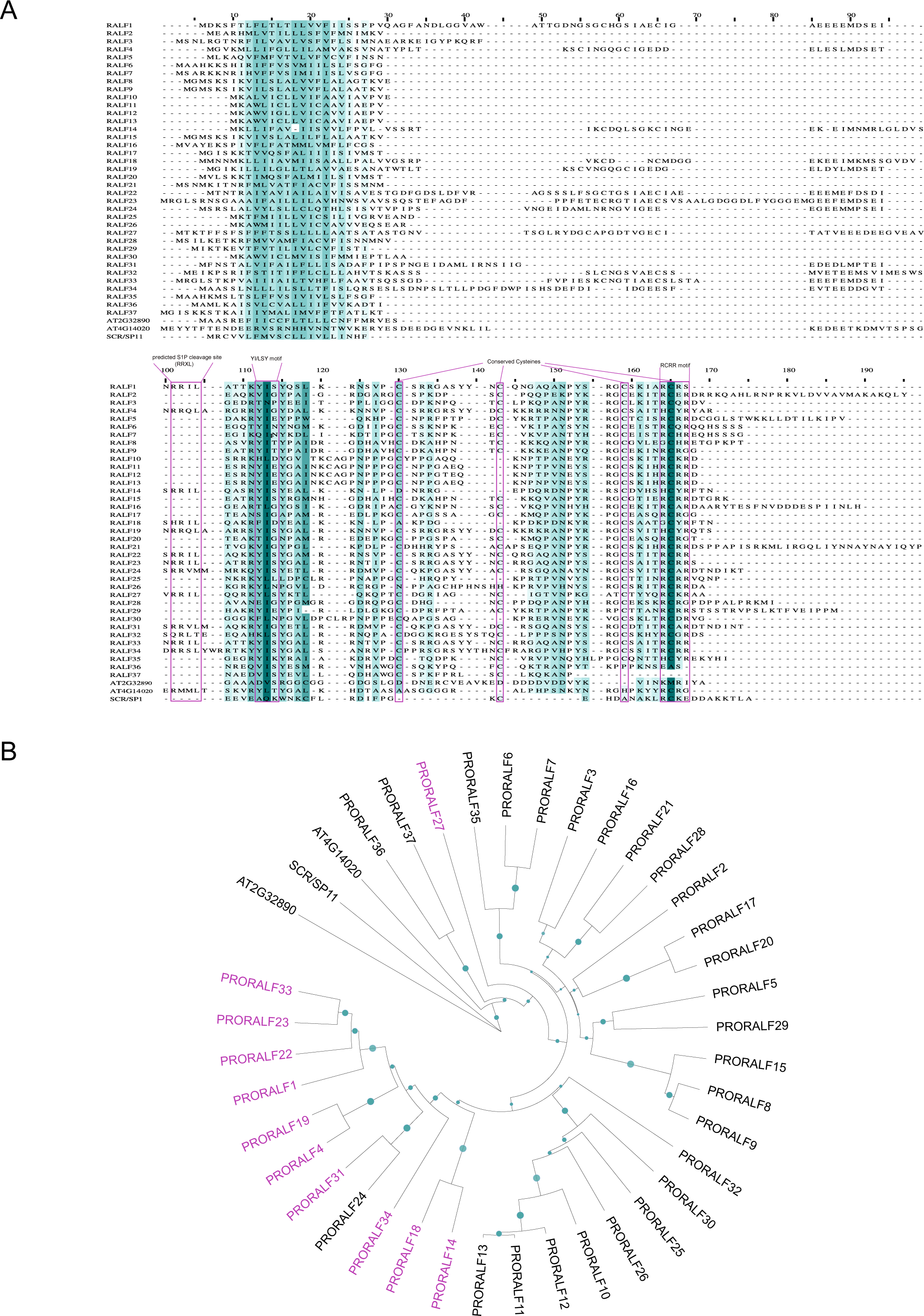
Re-evaluation of the *A. thaliana* RALF family. (A) Alignment of *At*PRORALFs, AT2G32890 and AT4G14020. Colour-code based on sequence conservation. The darker the colour, the more conserved the residue. Pink boxes indicate conserved motifs (B) Phylogenetic rooted tree of the *At*RALF peptides, AT2G32890 and AT4G14020. UPGMA tree inferred from the MUSCLE alignment displayed in Fig 1A. SCR/SP11 sequence was used to root the tree. RALFs highlighted in pink indicated those predicted to be cleaved by the protease S1P. Bootstrap values (1000 repetitions) above 50 % are represented by purple circles in the corresponding branches.

RALF17 is described as AT2G32890 in the aforementioned databases, while it corresponds to AT2G32885 in some publications (Cao and Shi, 2012; Morato do Canto et al., 2014; Stegmann et al., 2017). Protein sequence analysis however revealed that AT2G32890 lacks the YI/LSY motif including the conserved internal I/L residues (Fig 1A). AT2G32890 additionally lacks the conserved RCRR motif (Fig1A), and notably also misses the four cysteine residues that are positionally conserved across the RALF family (Fig 1A). Furthermore, the phylogenetic tree inferred from the alignment of RALF aminoacid sequences places AT2G32890 as an outgroup (Fig 1B). Altogether, we conclude that AT2G32890 has been mistakenly annotated as a RALF peptide.

When searching for AT2G32885 (proposed in some publications as RALF17) in TAIR10 and UniProt, we observed that this protein is annotated as RALF36. However, in other publications RALF36 corresponds to AT2G32785 (Morato do Canto et al., 2014; Stegmann et al., 2017). If AT2G32890 is not a RALF peptide as argued above, based on our phylogeny, we propose that AT2G32885 actually corresponds to RALF17. This is in agreement with previous studies (Morato do Canto et al., 2014; Stegmann et al., 2017). In contrast, we propose to assign AT2G32785 to RALF36 (Table I). It has part of the YI/LSY motif and the first cysteine bridge conserved, although it lacks the C-terminal RCRR motif (Fig 1A). Phylogenetic analysis indicates that it is a distant RALF clustering with RALF37 (Fig 1B). Furthermore, based on the information inferred from the alignment and the phylogenetic tree, it is difficult to conclude definitely whether RALF36 (AT2G32785) and RALF37 (AT2G32788) are genuine RALF peptides. As they present some of the important conserved residues, we nevertheless included them in our list of core RALF family members (Table I, Fig S1).

RALF35 has two gene identifiers associated with it: AT4G14020 and AT1G60913 (Haruta et al., 2014; Morato do Canto et al., 2014; Stegmann et al., 2017). Both have the highly conserved I/L amino acid within the YI/LSY motif as well as the two flanking tyrosine residues (Fig 1A). However, only AT1G60913 contains the four cysteine residues in the conserved positions; while AT4G14020 has only one cysteine at a conserved position. Moreover, AT4G14020 has part of the conserved RCRR motif, while AT1G60913 has not. Additionally, AT4G14020 lacks a predicted signal peptide and is thus likely non-secreted; it also does not cluster together with any other RALF in phylogenetic analysis (Fig 1B). Together, this indicates that AT4G14020 is not a bona-fide RALF peptide (Table I).

### Domain organization of the reannotated AtPRORALF family

Like other plant polypeptide hormones, *PRORALF* genes encode pre-pro-peptides of approximately 60-140 amino acids, which are predicted to undergo proteolytic processing to release bioactive RALF peptides (Fig 2; Olsson et al., 2019). PRORALFs have a N-terminal signal peptide for entry in the secretory pathway and the mature active peptide is located at the C-terminal part (Matos et al., 2008). Although 11 PRORALF proteins have a predicted subtilase cleavage site (RRXL, residues 101 to 104, Fig 1), so far only PRORALF23 and PRORALF22 have been experimentally shown to be cleaved by the subtilase SITE-1 PROTEASE (S1P) (Srivastava et al., 2009; Stegmann et al., 2017; Zhao et al., 2018). The protein domain organization of the PRORALFs that do not have the cleavage site suggests that they might not need to undergo subtilase-mediated proteolytic cleavage in order to release bioactive RALF peptides. Considering our proposed re-annotation of the *At*RALF family (Table 1), we depict the protein domain organization of the corresponding PRORALF proteins in Figure 2.

**Figure 2.**
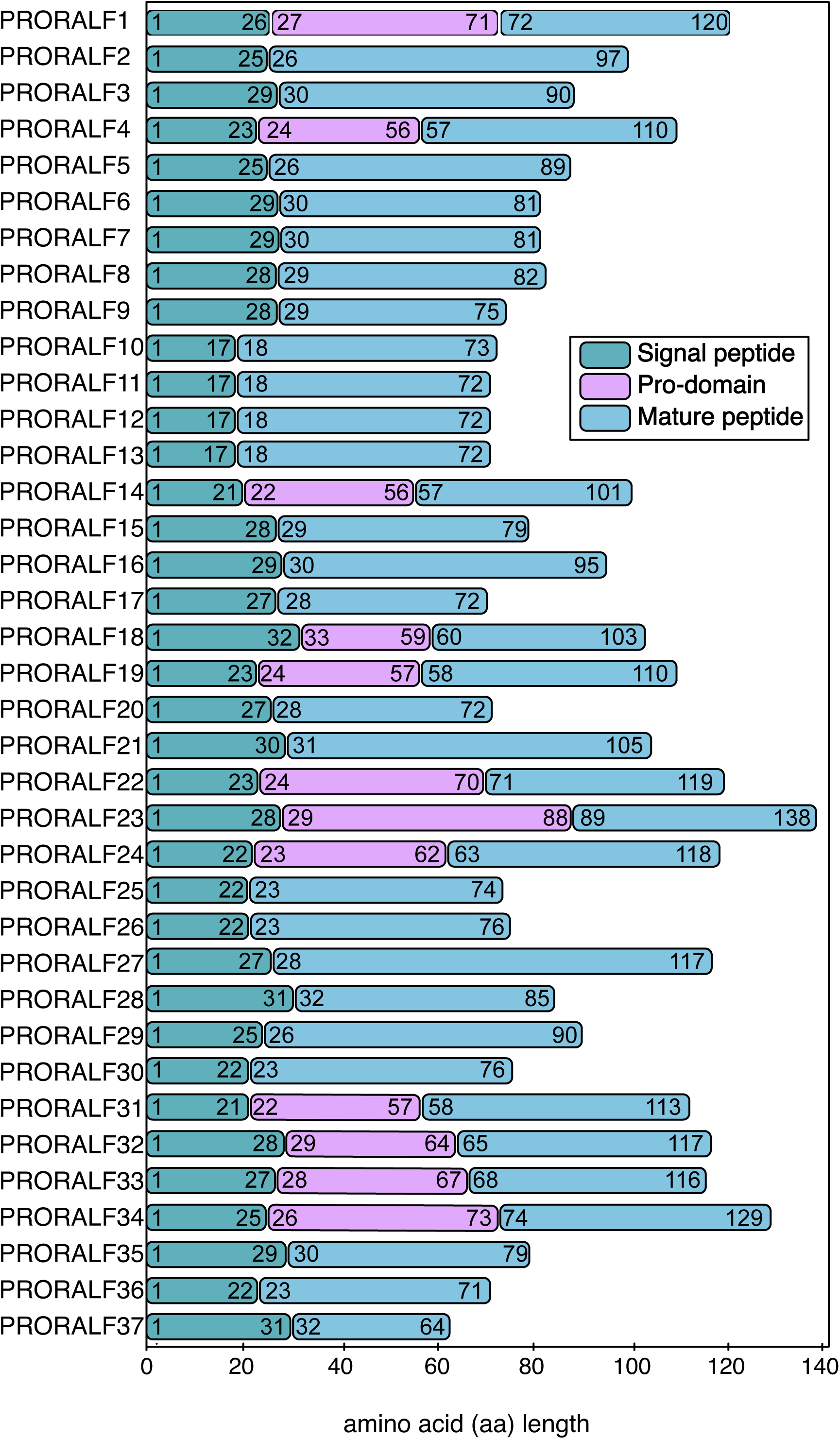
Schematic representation of *A. thaliana* PRORALFs domains. RALF peptides are expressed as protein precursors that require further processing steps.They contain an N-terminal SP (in green), variable pro domain for those which are predicted to be cleaved (pink) and a mature C-terminal part (blue).

### The majority of the *At*RALF peptides have growth inhibitory properties

One of the described functions of RALF peptides is to inhibit cell expansion and growth (Blackburn et al., 2020), but this is based on testing of only a few family members (Morato do Canto et al., 2014). Here, we screened 32 *At*RALF peptides for their bioactivity using seedling and root growth inhibition as read-outs (Fig 3). RALF11 and RALF12 were not synthesized because of their high sequence similarity with RALF13 (identical mature peptide), while RALF35, RALF36 and RALF37 could never be successfully synthesized, despite several attempts.

**Fig. 3.**
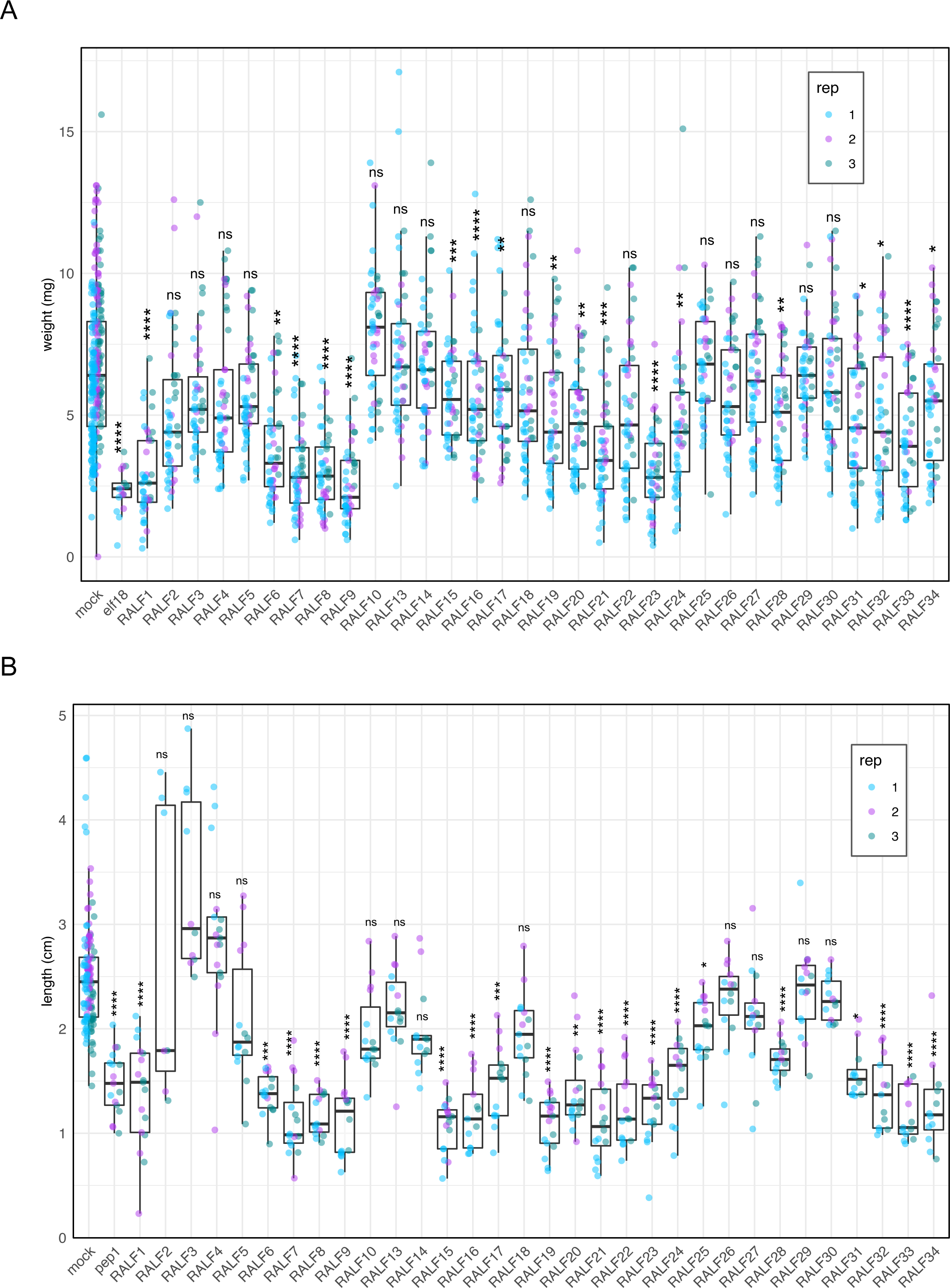
Effect of *At*RALF peptides on seedling and root growth inhibition. (A) Fresh weight of 12-day-old seedlings grown in the absence (mock) or presence of 1 μM RALF peptides (n=12) using elf18 (100 nM) as control. (B) Primary root length of 8-day-old seedlings grown in the absence (mock) or presence of 2 μM RALF peptides (n=6) using pep1 (10 nM) as control. Data from three independent experiments are shown (colours indicate different replicates). Upper and lower whiskers represent 1.5 times and −1.5 times interquartile range; upper and lower hinges represent 25% and 75% quartiles; middle represents median or 50% quartile. Asterisks indicate significance levels of a Kruskall-Wallis’s multiple comparison test, each treatment was compared to their corresponding mock: ns (p-value > 0.05), * (p-value ≤ 0.05), ** (p-value ≤ 0.01), *** (p-value ≤ 0.001) and **** (p-value ≤ 0.0001).

We treated 5-day-old seedlings with different synthetic RALF peptides (Table II) for 7 days before measuring seedling fresh weight or root length. We used the EF-Tu-derived peptide elf18 and the plant-derived peptide AtPep1 as positive controls for the seedling and root growth inhibition assays, respectively (Zipfel et al., 2006; Krol et al., 2010). Eighteen out of 32 (56 %) tested RALF peptides showed a significant seedling growth inhibition in the three independent biological experiments (Fig 3A).

**Table II.**
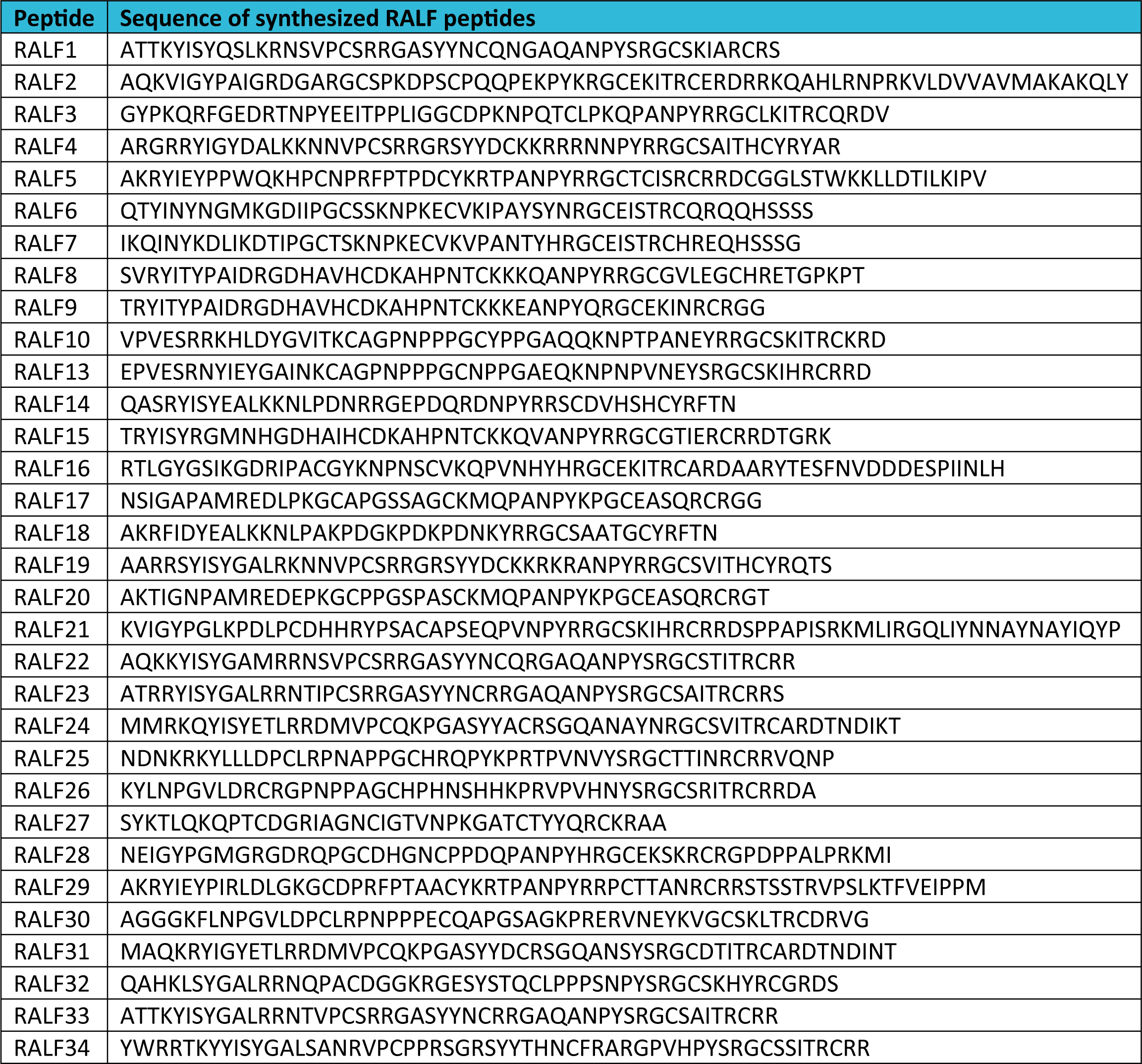
Sequences of RALF peptides synthesized. Aminoacidic sequence of the synthesized RALF peptides.

*AtRALF* genes and proposed receptor modules have a diverse expression patterns (Cao and Shi, 2012) and seedlings fresh weight is primarily determined by shoot biomass; it is therefore possible that no inhibition in the overall weight of the seedling may be observed if the receptor of a specific RALF peptide is only expressed in roots. For this reason, we also measured the primary root length after treatment with the different peptides. Strikingly, 20 out of 32 (63%) of the tested RALF peptides were also able to induce root growth inhibition (Fig 3B). The 18 RALF peptides that inhibited whole seedling growth were also able to inhibit root growth, while RALF22 and RALF25 only inhibited root growth. These data show that the majority of exogenously applied RALF peptides have the ability to inhibit growth under the conditions tested.

### The majority of growth-inhibitory *At*RALF peptides are FER-dependent

There are 17 CrRLK1L members in *A. thaliana* playing multiple and diverse roles, including cell growth, reproduction and responses to the environment (Franck et al., 2018; Blackburn et al., 2020). FER, the best characterized member of the family, is expressed throughout the plant, and has already been shown to mediate recognition of RALF1, RALF23 and RALF22 in diverse physiological contexts (Haruta et al., 2014; Stegmann et al., 2017; Zhao et al., 2018). As such, we performed seedling and root growth inhibition assays in the knock-out mutant line *fer-4* in comparison with Col-0 (Fig 4). Surprisingly, *fer-4* mutant seedlings were insensitive to 15 out of 18 (83 %) RALF peptides that inhibit seedling growth in Col-0 (Figs 3A and 4A). In the case of root growth inhibition, *fer-4* mutant seedlings were insensitive to 16 out of 20 (75 %) RALF peptides that inhibit root length in Col-0 (Figs 3B and 4B). The FER-independent RALF peptides that coincide between both assays are RALF28 and RALF34, which have the same effect in Col-0 and in the mutant line *fer-4* (Fig 4). In comparison, RALF20 is still able to mildly inhibit seedling growth in *fer-4* background but its root growth inhibitory effect depends on FER (Fig 4). Interestingly, RALF32 and RALF33 are FER-dependent in the seedling growth inhibition assay, but are still able to inhibit root length in the mutant line *fer-4* (Fig 4). Altogether, our data indicate that FER is involved in the perception and/or signalling pathway of the majority of the RALF peptides tested in the context of seedling and root growth inhibition.

**Fig. 4.**
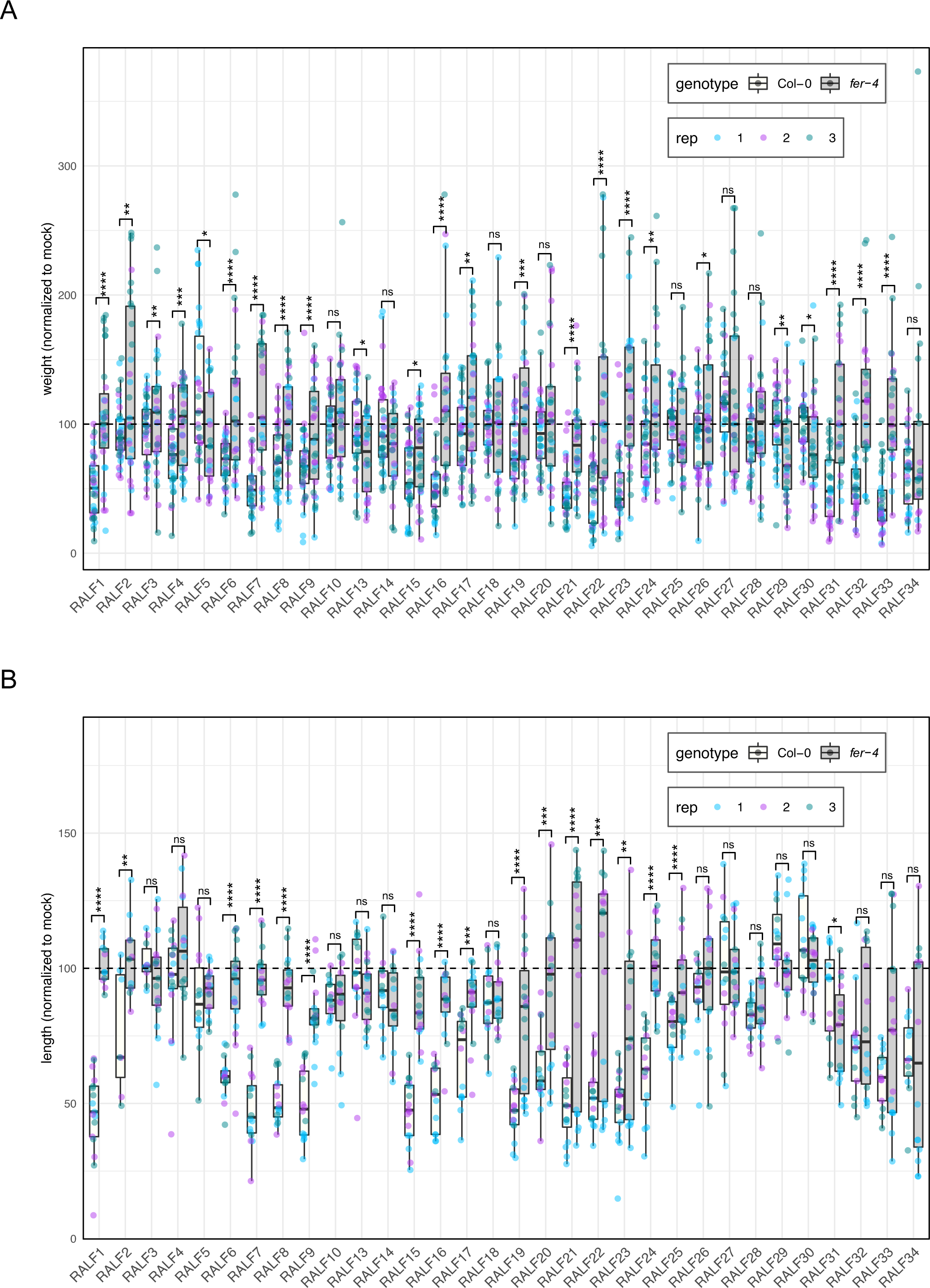
FER-dependency of *At*RALF peptides inducing growth inhibition. (A) Fresh weight of 12-day-old seedlings grown in the presence of 1 μM RALF peptides relative to mock (B) Primary root length of 8-day-old seedlings grown in the presence of 2 μM RALF peptides relative to mock. Mock primary root length of Col-0 (blue) and *fer-4* (pink) correspond to 100 % in the y axis. Data from three independent repetitions are shown (colours indicate different replicates). Upper and lower whiskers represent 1.5 times and −1.5 times interquartile range; upper and lower hinges represent 25% and 75% quartiles; middle represents median or 50% quartile. Asterisks indicate significance levels of a two-tailed T-test comparing each treatment in *fer-4* to the corresponding in Col-0: ns (p-value >0.05), * (p-value ≤ 0.05), ** (p-value ≤ 0.01), *** (p-value ≤ 0.001) and **** (p-value ≤ 0.0001).

## Discussion and Conclusions

The biological function of RALF peptides is an area of growing interest. Several studies on this family of peptides in different plant species have generated extensive knowledge about their roles and functions (Blackburn et al., 2020). Yet, even in the model plant *A. thaliana*, there is no agreement about the exact *At*RALF family composition (Olsen et al., 2002; Cao & Shi, 2012; Morato do Canto et al., 2014; Sharma et al., 2016; Campbell & Turner, 2017). Here, we gathered information from previous publications and databases to perform phylogenetic analyses of *At*RALF isoforms, and propose a new consensus annotation (Table I, Fig S2). By focusing on the model plant *A. thaliana*, we wanted to provide a more accurate annotation of the RALF peptide family rather than trying to provide evolutionary information on the family, as has been previously done (Cao & Shi, 2012; Sharma et al., 2016; Campbell & Turner, 2017). Based on previously identified conserved motifs (Pearce et al., 2001; Olsen et al., 2002; Matos et al., 2008; Pearce et al., 2010; Cao & Shi, 2012), we conclude that AT2G32890 and AT4G14020 are not genuine RALF peptides. AT2G32890 was probably annotated in the family as a mistake due to its close proximity on Chromosome 2 with the genuine RALF17 (AT2G32885). In turn, AT4G14020 was probably annotated in the family as a result of some degree of sequence similarity. It has been previously shown that tandem duplications played a dominant role in the evolution of *A. thaliana* RALF peptides (Cao & Shi, 2012; Campbell & Turner, 2017). It is possible that some of the newly duplicated genes have evolved under positive selection, causing changes in the protein sequence, which could explain why some of the proposed RALFs lack some of the conserved motifs. Future structural and physiological work, for example via protein chimeras, could investigate the biological relevance of the individual protein motifs as previously done for the conserved YI/LSY motif (Pearce et al., 2010).

Our results show that the majority of RALF peptides induce inhibitory effects on seedling weight and primary root length when exogenously applied (Fig 3). This result is consistent with a previous study that showed biological activity for nine recombinant RALF peptides (Morato do Canto et al., 2014). Notably, despite being closely related and varying only in seven amino acids, RALF19, but not RALF4, induced growth inhibition (Fig 3). RALF19 and RALF4 were previously tested for alkalinisation activity, which RALF19 possesses but RALF4 does not (Morato do Canto et al., 2014). This might indicate that the growth inhibition activity is linked to the ability of these peptides to increase the pH of the extracellular space. Additionally, treatment with RALF1 suppresses cell elongation of the primary root by activating FER, which in turns causes the phosphorylation of the plasma membrane H^+^-ADENOSINE TRIPHOSPHATASE 2 (AHA2), which inhibits proton transport (Haruta et al., 2014). Whether this pathway is transferable to the rest of the family remains elusive. Future work expanded to the whole family should determine this unclear link between pH and growth.

FER is the best studied member of the CrRLK1L family, and has been shown to be involved in numerous physiological processes (Franck et al., 2018; Blackburn et al., 2020). FER was recently shown to recognise diverse RALFs, such as RALF1, RALF22 and RALF23 (Haruta et al., 2014; Stegmann et al., 2017; Zhao et al., 2018; Xiao et al., 2019). Here, we show that FER is required for the inhibitory activity of the majority of RALF peptides in the context of growth inhibition (Fig 4). FER is widely expressed throughout the plant (Lindner et al., 2012). As our assays rely on the exogenous treatment with synthetic RALF peptides, our results do not necessarily imply that FER is the receptor for the corresponding endogenous RALF peptides. Nevertheless, it is interesting that not all growth-inhibitory RALF peptides are FER-dependent, which would argue for some level of specificity. Additionally, some RALF peptides, such as RALF32 and RALF33, are FER-dependent when inhibiting seedling growth but are still able to inhibit root growth in the *fer-4* mutant line (Fig 4). This suggests that these RALF peptides might be perceived by different receptor complexes in the root and in the shoot. Also, it is notable that different CrRLK1Ls have recently been shown to work together as part of hetero-multimeric protein complexes to mediate RALF perception or control most likely RALF-regulated processes. For example, RALF4 and 19 are proposed to be perceived by a complex involving ANX1/2 and BUPS1/2 controlling pollen tube growth and integrity (Ge et al., 2017), while ANJEA (ANJ) and HERCULES RECEPTOR KINASE 1 (HERK1) regulate pollen tube reception (Galindo-Trigo et al., 2020). Also, while THE1 is the receptor for RALF34, *fer-4* is also affected in RALF34 responsiveness (Gonneau et al., 2018), suggesting that THE1 and FER might form a heteromeric complex to control responsiveness to cellulose biosynthesis inhibition. In our hands, however, RALF34-induced growth inhibition was similar in Col-0 and *fer-4* mutant background (Fig 4). This suggests that, while THE1 is the primary RALF34 receptor, it might form distinct heteromeric complexes with different CrRLK1L-family members depending on the context.

Our results provide the basis for the future identification of RALF-CrRLK1L ligand-receptor pairs. It will however be essential to determine overlapping expression patterns of different *PRORALF* and *CrRLK1L* genes across different organs, tissues and cell types, and during different developmental stages using either transcriptional reporters (Gonneau et al., 2018) or capitalizing on recent quantitative proteomics studies of the *A. thaliana* proteome (Zhang et al., 2019; Mergner et al., 2020; Bassal et al., 2020). These approaches will guide downstream biochemical/biophysical characterizations of ligand-receptor binding and potential heteromeric CrRLK1L complexes, as well as the genetic characterization of *PRORALF* and *CrRLK1L* genes, which otherwise can suffer from functional redundancy and pleiotropic issues. For example, recently-developed approaches such as cell-specific CRISPR/Cas9-mediated genome editing could be used to generate high-order receptor and ligand mutants (Decaestecker et al., 2019; Wang et al., 2019). Together, these integrated approaches will be needed to decipher the complex signalling network that RALF peptides and their corresponding receptors weave in their native contexts.

## Material and Methods

### Plant growth and conditions

*Arabidopsis thaliana* seeds were surface-sterilized using 70 % and 100 % ethanol for 20 minutes and grown on 1/2 Murashige and Skoog (MS) media with 1% sucrose, adjusted to pH 5.8 using KOH, with or without 0.9 % agar at 20 °C and a 16h-photoperiod. The *fer-4* seeds were kindly provided by Alice Cheung (University of Massachusetts).

### Peptides

RALF peptides were synthesized (Table II) by SciLight Biotechnology LLC (www.scilight-peptide.com) with a purity of >85 %. All peptides were dissolved in sterile pure water for usage and stored at −20 °C at a concentration of 1 mM.

### Phylogenetic analysis

Multiple sequence alignments of the full-length or mature peptide sequences were created using the MUSCLE algorithm with the MEGA X software (Kumar et al., 2018). Sequence alignment was coloured according to sequence conservation and amino acid type using the software Jalview. Phylogenetic rooted trees were constructed with the MEGA X software by using the UPGMA algorithm with the default parameters. Bootstrapping was performed 1000 times. The inferred trees were visualized using iTOL (https://itol.embl.de/) (Letunic and Bork, 2019).

### Seedling growth inhibition assay

Seeds were surface-sterilized and grown on MS agar plates for 5 days before transferring individual seedlings in each well of a 48-well plate containing 500 μL per well of MS medium containing 1 μM RALF, or 5 nM elf18 as control. Seedlings weight was measured 7 days later. Control seedlings were grown under identical conditions in a peptide-free medium. The experiments were repeated 3 times using independent biological replicates. Twelve seedlings for each treatment were measured.

### Root growth inhibition assay

Seeds were surface-sterilized and vertically grown on MS agar plates for 5 days before transferring 6 seedlings to each well of a 12-well plate containing 4 mL per well of MS medium containing 2 μM RALF using 10 nM AtPep1 as control. Seedlings were transferred 3 days later to solid MS plates. Control seedlings were grown under identical conditions in a peptide-free medium. Primary root length was measured by scanning the plates and quantified using the software Fiji (https://imagej.net/Fiji) (Schindelin et al., 2012). Experiments were repeated 3 times using independent biological replicates. Roots from approximately 6 seedlings for each treatment and genotype were measured.

### Statistical analysis

Statistical analysis was performed applying non-parametric Kruskal-Wallis multiple comparison test, comparing every treatment to its respective mock control using Prism 8.0 (GraphPad Software). Similar significance levels were obtained when transforming the data to normal distribution and performing a Two-way ANOVA test followed by Dunnett’s post-hoc test using the software R. Two-tailed t-tests were performed to assess the significant differences between Col-0 and *fer-4* in growth inhibition experiments, using Prism 8.0 (GraphPad Software).

## Acknowledgments

We thank all the members of the Zipfel group, and most particularly Marta Bjornson and Isabel Monte, for fruitful discussions and guidance. We thank Joop Vermeer and Vinay Shekhar for providing access to the plate scanner. This work was supported by the European Research Council under the Grant Agreement no. 773153 (grant ‘IMMUNO-PEPTALK’), the University of Zürich, and the Swiss National Science Foundation grant no. 31003A_182625. C.M.F. is supported by a post-doctoral fellowship (EMBO ALTF 512-2019) from the European Molecular Biology Organization. Authors have no competing interests.

All data is available in the manuscript or the supplementary materials.

